# A Case Study: Using Passive Acoustic Monitoring Devices In A Growing Urban Landscape To Monitor Avian Diversity

**DOI:** 10.1101/2025.07.21.665881

**Authors:** Anshul Gupta, CS Swathi, Asher Jesudoss, Anusha Shankar, Harsha Kumar

## Abstract

With the increase of urbanization around the world, key habitats such as grasslands and scrublands are disappearing, posing significant threats to species that rely on such habitats. Passive acoustic monitoring (PAM) has started to gain interest as a reliable tool for biodiversity monitoring. This study assesses whether PAM can be an effective way to monitor biodiversity with relatively low effort when compared to traditional survey methods. Using PAM, a mixed grassland and scrubland ecosystem was monitored for five months for avifaunal diversity. Data that was collected from dusk to dawn using audio recorders deployed on the study site were run through BirdNET analyzer under default detectors. Top detections for each species were compared with avian vocalization libraries manually. Out of 135 species detected, 76 were true positives, resulting in a species identification accuracy of 56.3%. Notably, among the species that were true detections, 21 species were found to have confidence scores larger than 0.99. In addition, 15 of the detected species were found to be migratory in this area, and three were found to be rare. This study demonstrates how PAM can be used to monitor biodiversity in a species-rich but understudied area such as Southern India, to identify both cryptic and nocturnal species that might be omitted from standard field surveys.

## Introduction

In recent years, urbanization has increased rapidly around the world, changing landscapes and promoting the urban heat island effect, leading to increased temperatures in regions with dense infrastructure.^1^ Anthropogenic development creates some of the greatest local extinction rates and eliminates the majority of local native species.^2^ While grasslands and scrublands cover almost 60% of the world’s land area, they are being increasingly used for new development projects in cities.^3–5^ This poses a significant threat to critically endangered species that rely on these habitats, such as the Great Indian Bustard (*Ardeotis nigriceps*) and the Speckled Ground Squirrel (*Spermophilus suslicus*).^6,7^

Studies have already documented a decrease in species richness of urban areas over time.^8^ Therefore, it is important to monitor such fragile landscapes and conduct ecological surveys as part of urban and conservation planning. One group of organisms that has been drastically affected by urbanization is birds.^9,10^ Birds are important bioindicators of their environment and have previously been used to assess the health of scrublands and as indicators of pollution in aquatic and terrestrial environments.^11,12^ They offer ecological services such as pollination, seed dispersal, and pest management.^13,14^ Studying these organisms can thus be an important contribution to the study of the ecosystem as a whole.

Passive acoustic monitoring (PAM) is an observational tool used to study ecology without interfering with the environment. Research interest in PAM is rapidly growing: the hardware has become more compact and affordable, and audio analysis software is becoming more sophisticated.^15^ Apart from this, PAM has been shown to perform as well as or better than observer-based monitoring.^16^ Within the past decade, PAM has been used in studies ranging from estimating primate occupancy to finding critically endangered species such as the Jerdon’s Courser (*Rhinoptilus bitorquatus*), proving its value in field monitoring in a range of environments, including several applications for conservation decisions.^17,18^

Our goal was to conduct an ecological assessment of scrub-grassland avian species richness in a changing urban landscape. To achieve this, we used BirdNET analyzer with default detectors to identify possible species present on the study site.^19^ Next, we qualitatively (audio-visually) inspected top detections to confirm the presence of the species. We consulted expert naturalists to validate any ambiguous calls. An implicit goal of the project was to quantify the accuracy of default detectors for species in Southern India that are poorly studied. Passive acoustic monitoring can therefore be a powerful tool to efficiently collect data on species’ occurrences across large areas, especially in urban landscapes that are biodiverse and data-deficient.

## Methodology

### A. Study site

Tata Institute of Fundamental Research Hyderabad (TIFRH) campus is a 200-acre mixed habitat ecosystem, in Gopanpally, Hyderabad, Telangana, India. The TIFRH campus was partitioned from the University of Hyderabad in 2017, whose bird diversity has been well documented.^20,21^ The site has two blocks of scrub-grassland habitat with rocky outcrops that can be characterized predominantly by Persimmons (*Diospyros spp*.), Mountain Pomegranate (*Catunaregam spinosa*), Indian Jasmine (*Jasminum auriculatum*), Mysore Sumac (*Searsia mysorensis*), Neem (*Azadirachta indica*), and Golden Shower Tree (*Cassia fistula*) communities. All together, this land supports around 80 species of birds and other fauna such as the Spotted Deer (*Axis axis*), and the vulnerable Indian Star Tortoise (*Geochelone elegans*) as seen by the authors of this study.^22^

### B. Acoustic Sampling

Seventeen AudioMoths (audio recorders) were set up equally spaced across the site, fastened to trees two meters above the ground.^23^ They each recorded for an average of 25 days in Nov-Dec, 25 days in Jan-Feb, and 13 days in March between 5:30 p.m. and 6:30 a.m. each day. The first 20 seconds of each minute were sampled, given that temporal stratification of sampling at shorter timescales (20 seconds at the start of every minute compared to 20 minutes at the start of every hour) is more useful for measures of both alpha and gamma diversity.^24,25^ This resulted in a total of approximately 3,900 recording hours.

### C. Data Analysis

Given the volume of the data, we used the BirdNET (Model version 2.4, GUI version 2.0.0) analyzer with custom settings to annotate data, as outlined below. Then, we manually validated the results using Raven Pro 1.6.5 to ascertain the presence of each species.^26,27^ Lastly, the accuracy of BirdNET analyzer was quantitatively assessed.

#### (i) BirdNET detections

All recordings were run through the BirdNET analyzer developed by the K. Lisa Yang Center for Conservation Bioacoustics to identify the bird species that might be present in them.^26^ A confidence threshold (confidence of bird occupancy estimated by the detector) of 0.75 was used, as anything lower than 0.75 could have resulted in false positives. The geo-location filter was set to the coordinates of the neighbouring University of Hyderabad (17.454654, 78.328636). The location filter threshold (minimum occurrence probability for a species to be included) was set to 0.01 to include rare or vagrant species that could be found in and around the study site. Outputs were saved as Raven selection tables and combined to facilitate further inspection. All other settings were left at default.

#### (ii) Validating birdNET detections

To validate whether the detector detected species that were actually present (true positives), we followed a custom pipeline (Figure 2). After selecting a species from detections, we first checked the eBird records for Telangana, India. If the species had not been reported before, it was considered a misidentification. If it had been reported before, the detections were sorted from highest to lowest confidence. The top 10 detections with the highest confidences were selected for review. If the bird was found confidently by matching it with recordings from Macaulay Library, it was considered a true detection.^28^ Ambiguous calls that could not be annotated through audio-visual inspection of spectrograms were sorted into correct or incorrect identifications by expert naturalists. In such cases, the detections may not necessarily have a Macaulay Library ID link because confirmations were based on the experts’ acquired knowledge.^29^

**Figure 1.**
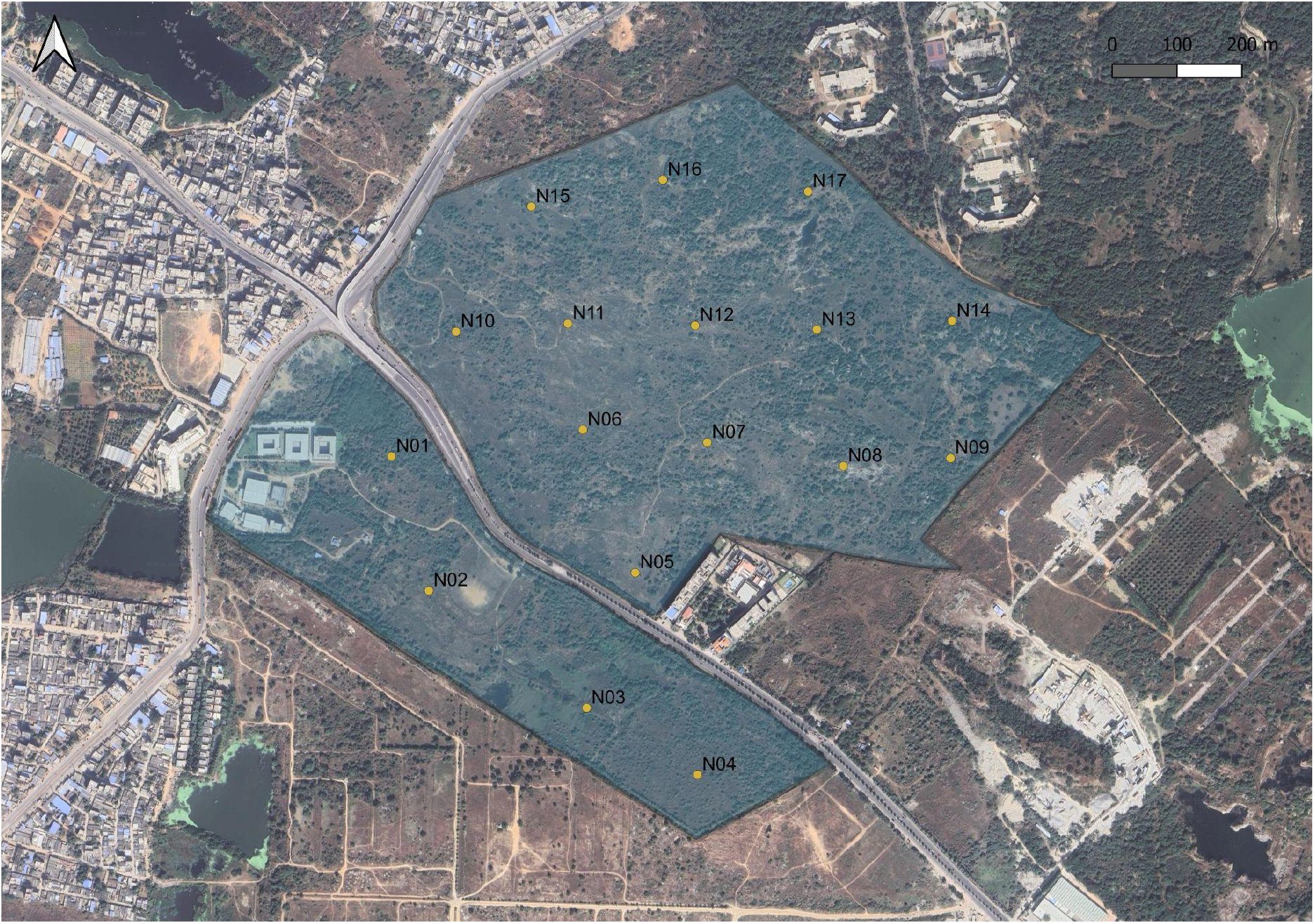
Map of points where AudioMoths were located (17.4456677, 78.3116856)

**Figure 2.**
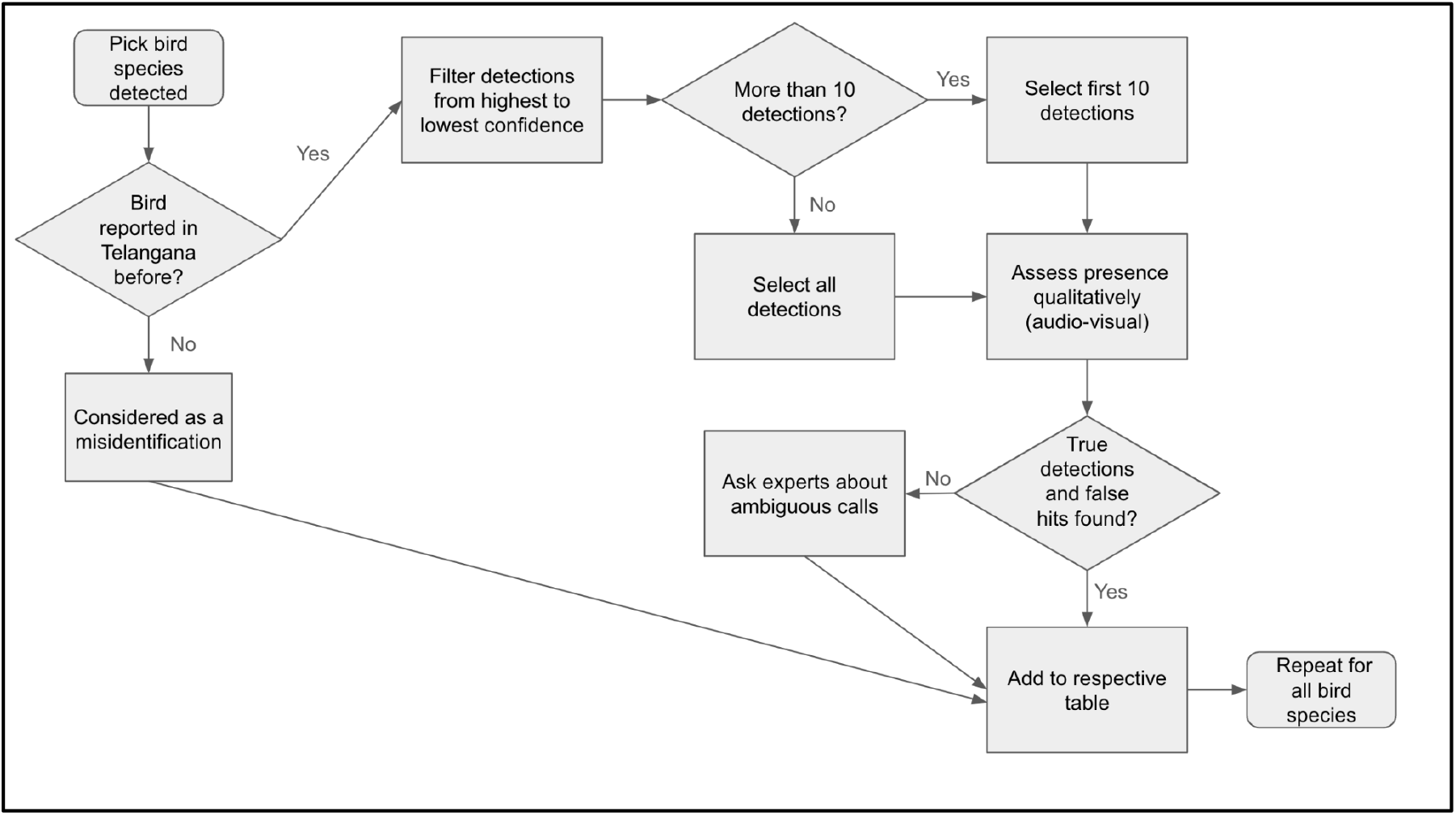
Flowchart for validating BirdNET detections

**Figure 3.**
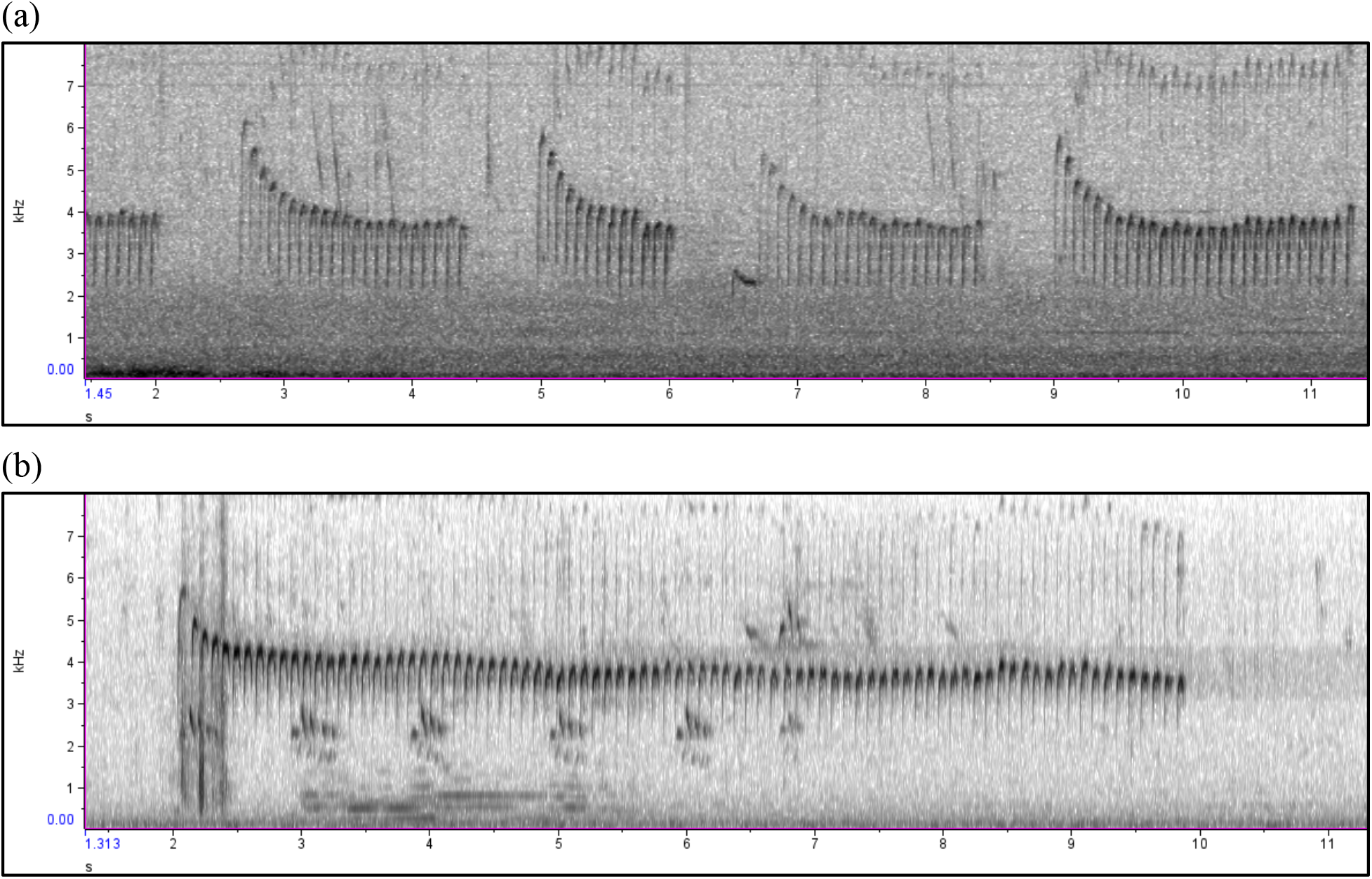
Spectrograms of the Common Babbler (*Argya caudata*). Spectrogram (a) was recorded on March 16, 2025, at 06:25 a.m. from Recorder 12 on the study site. Spectrogram (b) is a reference call from Xeno-canto.^35^ The song consists of sharp, quick notes that start at a higher frequency (around 6 kHz) and decrease until they remain stable (at around 4 kHz). As seen in spectrograms (a) and (b), this is a common feature. Length of vocalizations produced by this species is variable, but other spectral features are relatively invariant.

**Figure 4.**
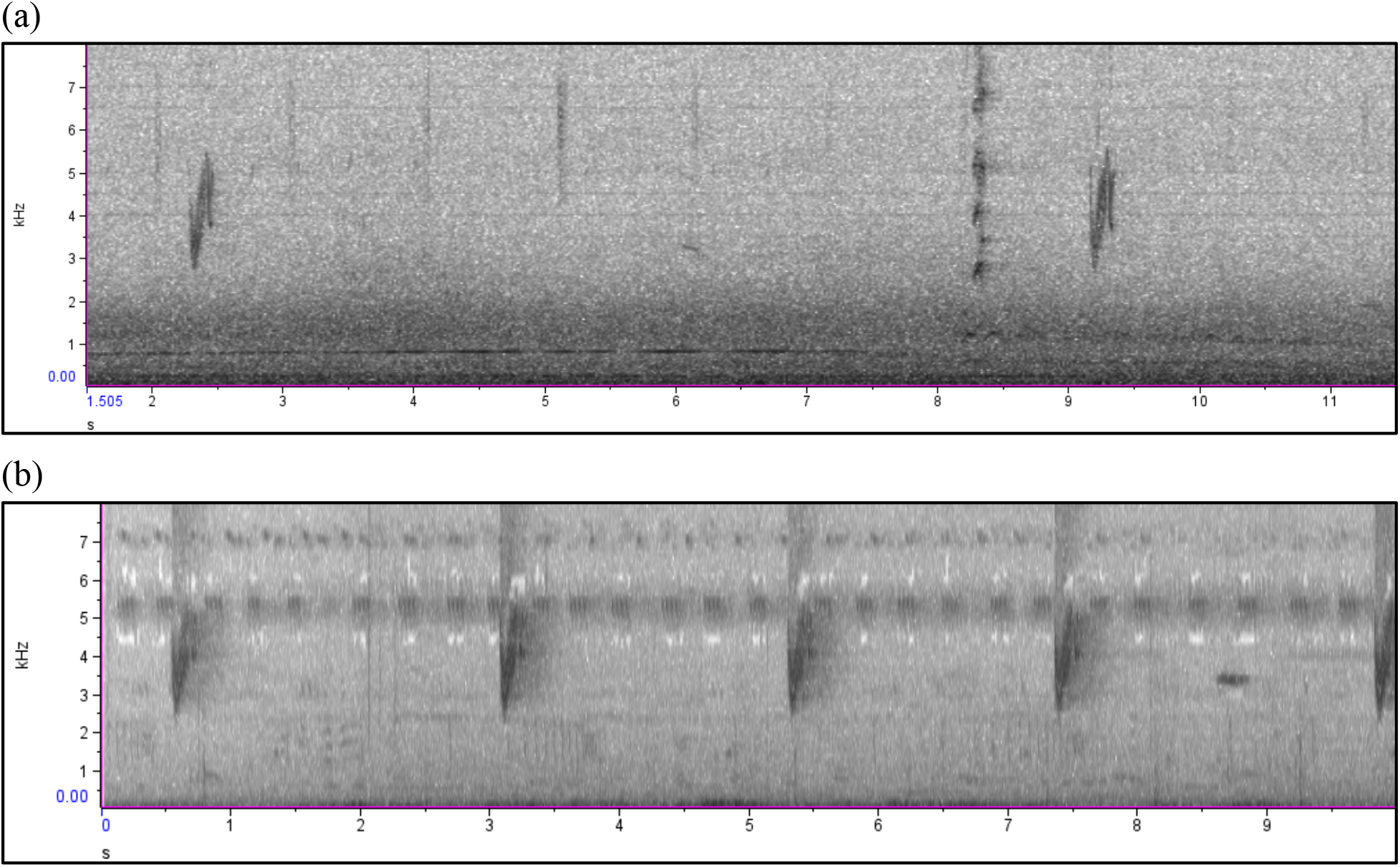
Spectrograms of the Savanna Nightjar (*Caprimulgus affinis*). Spectrogram (a) was recorded on March 27, 2025, at 04:08 a.m. from Recorder 04 on the study site. Spectrogram (b) is a reference call from Xeno-canto.^36^ The calls are a quick down-up chirp (at around 3 kHz to 5kHz). As seen in spectrograms (a) and (b), this is a common feature.

This process was repeated for every detected species. If the species was found in one set of data (for example, Nov-Dec), it was not checked again in any of the other months.

#### (iii) Assessing the accuracy of BirdNET analyzer detectors

Using the data from the tables from section C(ii), we calculated a percentage accuracy for default BirdNET detectors. This was calculated using the formula:

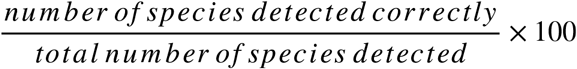

To find the percentage of incorrect detections, the value from the above formula was subtracted from 100.

## Results

A total of 135 species were detected, out of which 76 were correctly identified and 59 were misidentified. Of these species, expert validation was needed for nine species to confirm presence. While all of the species positively detected were listed as ‘Least Concern’ in the IUCN Red List, 15 species are known to be migratory.^30,31^ The percentage accuracy of the detector was estimated to be 56.3% as calculated based on the formula in C(iii) of the methodology.

### A. Species detected and found in data

While 21 species were detected with a confidence of above 0.99, nine of those species were detected with a confidence of 0.999 or above (Table 1). These species were namely the Gray Francolin (*Ortygornis pondicerianus*), Painted Francolin (*Francolinus pictus*), Indian Nightjar (*Caprimulgus asiaticus*), Indian Thick-knee (*Burhinus indicus*), Yellow-wattled Lapwing (*Vanellus malabaricus*), Red-wattled Lapwing (*Vanellus indicus*), Blue-tailed Bee-eater (*Merops philippinus*), Yellow-billed Babbler (*Argya affinis*), and Common Babbler (*Argya caudata*). For these species, the default detectors on BirdNET could potentially be used to monitor them for presence-absence.

**Table 1.**
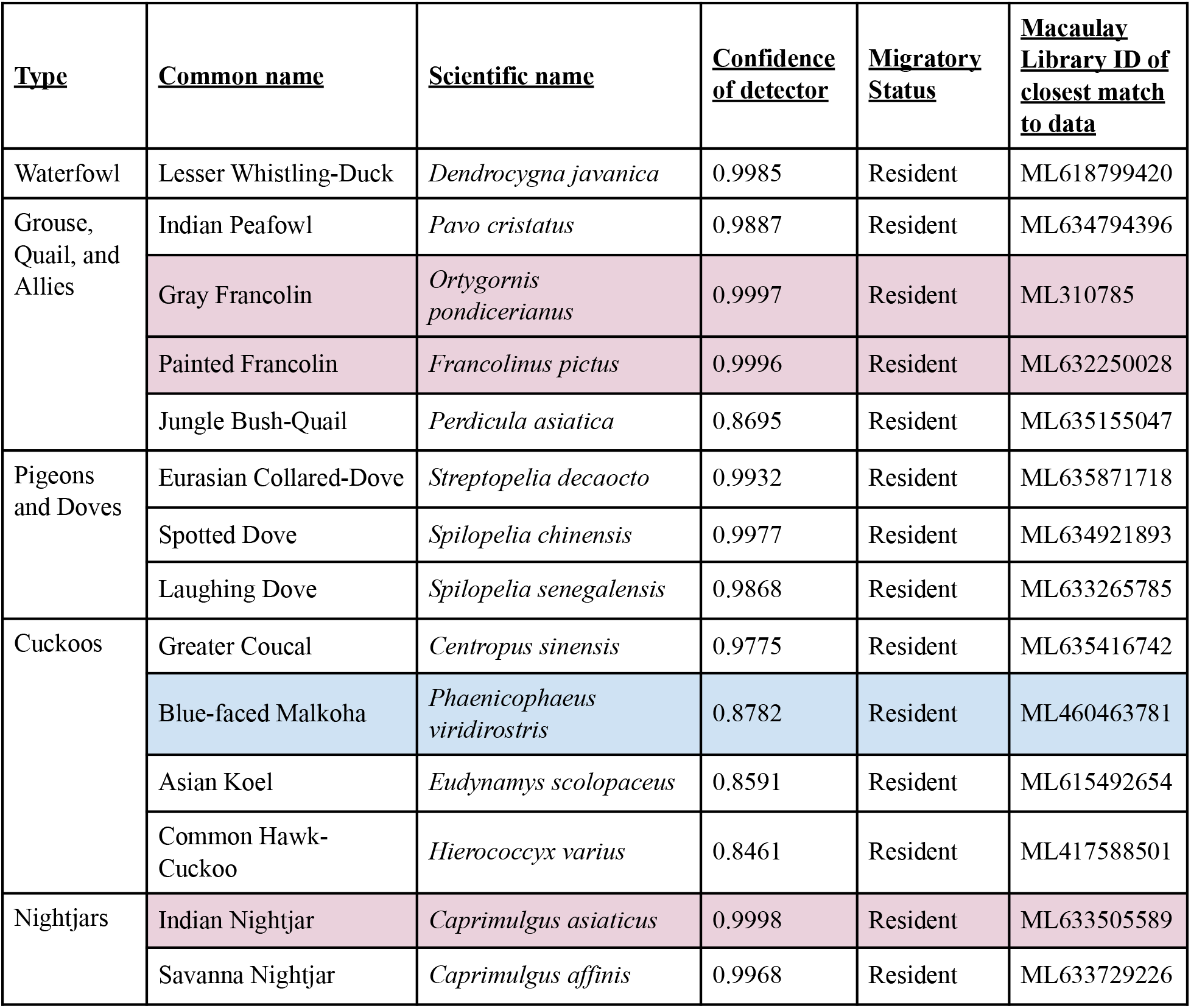
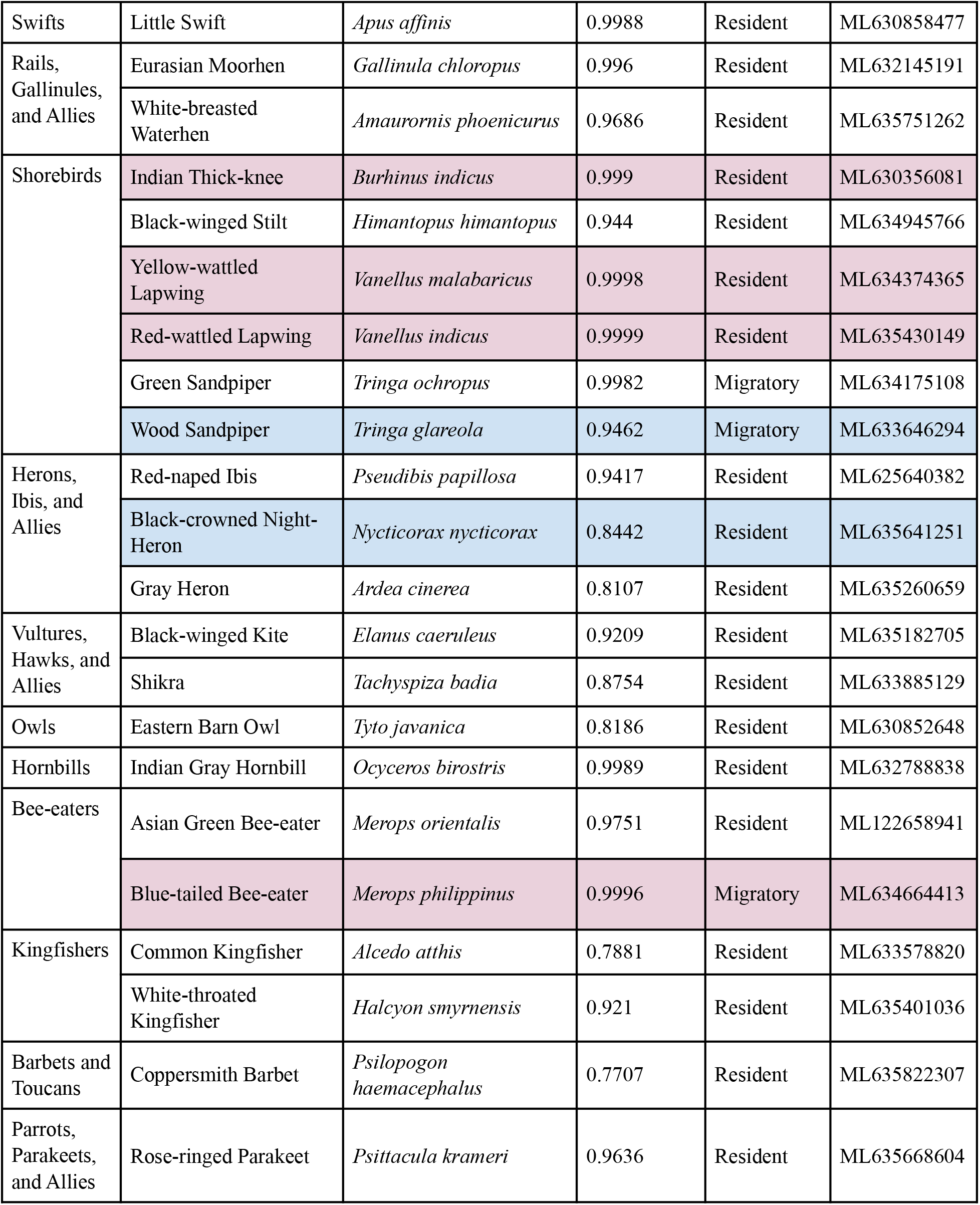
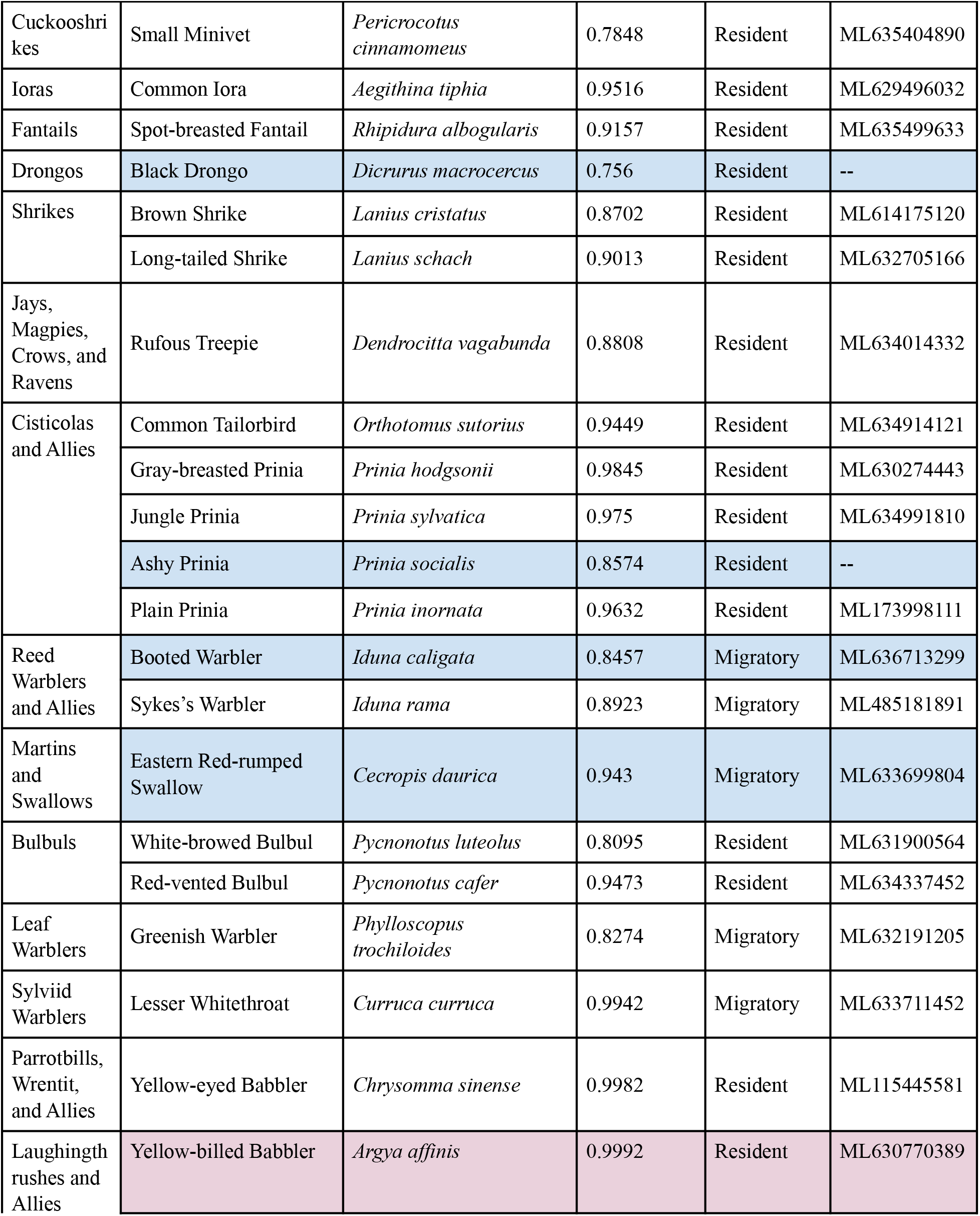
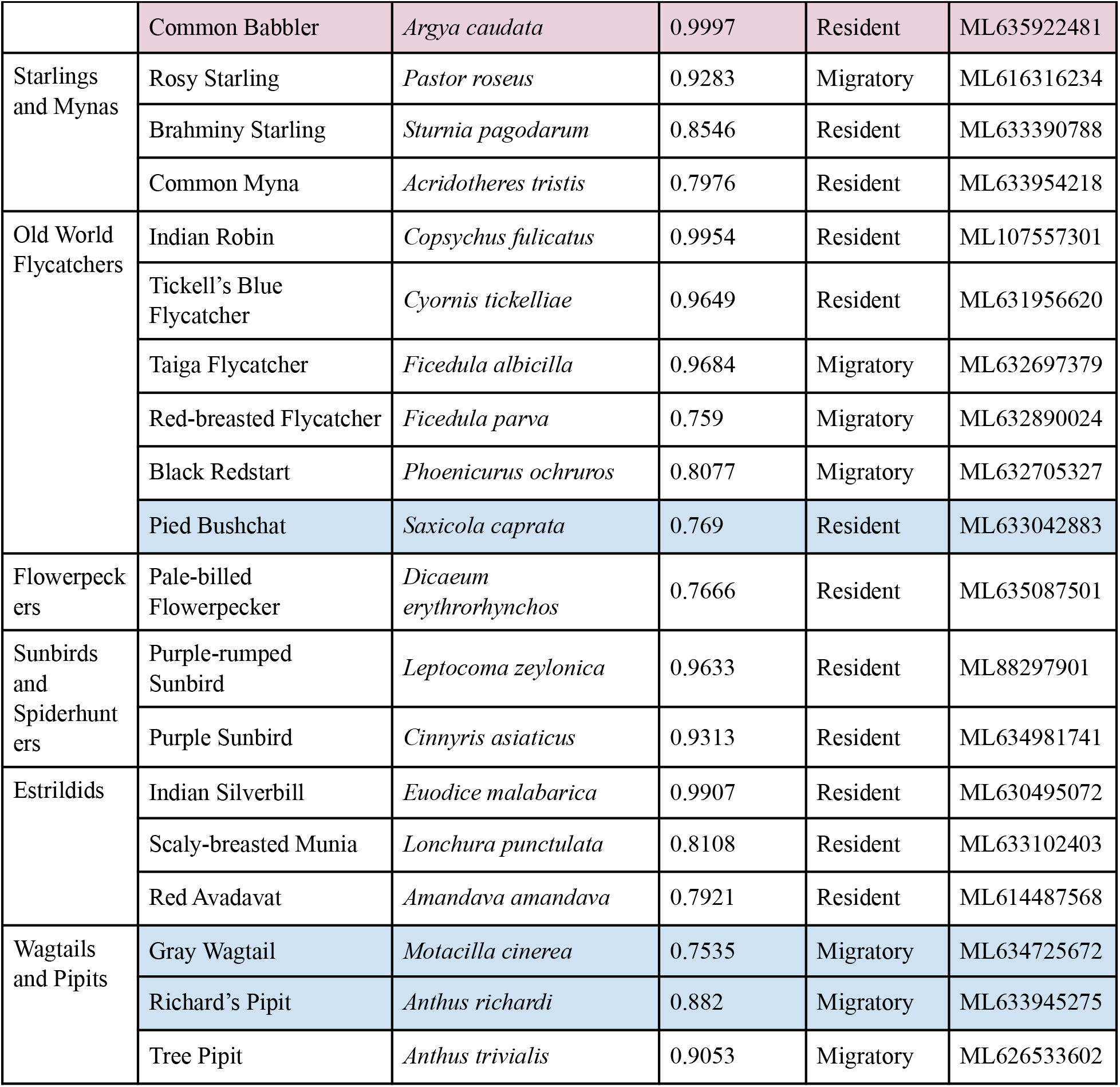
List of species with a confirmed presence from the dataset sorted by family. Confidence of BirdNET detections, migratory and IUCN status of species, in addition to species validated by experts, are shown above.^28,33^ Rows in blue are validated by experts. Rows in pink are species that can be detected effectively using default BirdNET detectors.

### B. Spectrograms of some of the rare species found

The spectrograms from data collected are shown for two rare species along with matches from Xeno-canto, a digital repository of animal sounds, below.^34^ Our detections included rare species such as the Common Babbler (*Argya caudata*) that has not been reported on the eBird hotspot for the study site.^22^The species was last reported at the University of Hyderabad, the nearest well-documented site, on 12 August 1992.^21^ While the species hasn’t been reported on eBird since, it was seen on the study site in 2020 by an expert independently. We detected calls of the species on November 16, 2024, and March 16, 2025.

Another rare species we detected on our study site included the Savanna Nightjar (*Caprimulgus affinis*), which has not been reported on the eBird hotspot for the study site nor the University of Hyderabad.^21,22^ Only two calls of this bird were detected in the entire dataset, and both were from the same 20-second recording. This suggests that it was a temporary visitor.

### C. Species misidentified in the data

A total of 59 species were misidentified, including 3 species with no records in the state of Telangana. Of these 59 species that were false detections, 15 were detected with a confidence of above 0.95, with 1 species, the Black-tailed Godwit (*Limosa limosa*), being detected with a confidence of above 0.99 (Supplementary Table 1). For these species, better detectors need to be built for PAM.

## Discussion

This study focuses on assessing growing urban landscapes for avifaunal biodiversity using passive acoustic monitoring (PAM). Biodiversity assessment and monitoring is often important in the contexts of city planning and environmental protection policies that are based around such primary data. We collected around 3,900 recorder-hours of data across 17 recorders over a 200-acre campus in the Hyderabad area in Southern India from November 2024 to March 2025. We used PAM to assess how effective this method is in identifying species in the region. The default BirdNET detectors identified 135 species, but 59 were misidentified, while 56.3% (76 species) were correctly identified. Of the 76 species positively detected, 19.7% (15 species) were migratory and 11.8% (nine species) needed expert validation to confirm the identity from the detector. PAM is therefore becoming a promising tool to assess biodiversity in the tropics. We simply need more training data to improve the detectors.

Although the calculated accuracy of 56.3% is low, it is similar to expected values shown by Merlin Sound ID for this geo-location.^37^ Compared to places such as the U.S.A. and Australia, much less sound recording data are available for birds in South India.^38^ For 21 species on our study site, the detection probabilities were above 0.99, given that their unique vocalizations did not overlap with other biotic, abiotic, or anthropogenic sounds. The Painted Francolin and the Indian Nightjar have distinct vocalizations that were picked up correctly with a high confidence of greater than 0.999. For these species, passive acoustic monitoring using default BirdNET detectors is viable in a variety of contexts (Table 1). Additionally, birds that are hard to distinguish visually can be further mapped using these methods. The Richard’s Pipit is a cryptic species that was found on the study site, but is often misidentified as other common pipits on eBird.^39^ Using vocalizations can be particularly useful for such species since the vocalizations are distinct and can potentially be used to map distributions.

This study serves as an example of the potential PAM offers for finding rare and migratory species across large areas of mixed scrubland and grassland habitats that would otherwise be difficult to cover with traditional survey methods. Some of the rare species found during this study were the Common Babbler (*Argya caudata*), Savanna Nightjar (*Caprimulgus affinis*), and Richard’s Pipit (*Anthus richardi*), the latter two of which are new records for the study site and have never been reported at the University of Hyderabad.^21,22^ Out of the 15 migratory species found, seven species, which include the Green Sandpiper (*Tringa ochropus*), Blue-tailed Bee-eater (*Merops philippinus*), Taiga Flycatcher (*Ficedula albicilla*), Red-breasted Flycatcher (*Ficedula parva*), Black Redstart (*Phoenicurus ochruros*), Gray Wagtail (*Motacilla cinerea*), and Tree Pipit (*Anthus trivialis*), are uncommon for the area. While these are interesting findings, our study is not without limitations.

For 15 species, the detector misidentified a variety of sounds as birds with a high confidence of above 0.95. Nocturnal insects were misidentified as diurnal birds such as the Ashy Prinia and the Plain Prinia, which use vocalizations at similar frequencies. For such misidentified species in South India, better detectors that rely on large training datasets are needed. While the presence of the Common Babbler was not recorded visually, we cannot account for other species mimicking them. For example, the Long-tailed Shrike (*Lanius schach*), Black Drongo (*Dicrurus macrocercus*), and Ashy Woodswallow (*Artamus fuscus*) are known to mimic a variety of calls.^40–42^ Since the Common Babbler has not been seen in the area since 1992 (and briefly in 2020), it is possible that the bird was not present on the site at the time of recording and that a different species was mimicking its call. Given that cultural transmission of learnt vocalizations is very strong in birds representing these genera, a visual confirmation of the Common Babbler would be necessary to confirm its presence.^43,44^ For a species like the Savanna Nightjar, however, there are no species that are known to mimic their calls. It therefore seems reasonable to conclude that the Savanna Nightjar was present on the study site.

We used a combination of different methods such as the deployment of audio recorders, use of BirdNET analyzer to detect vocalizations of species, and expert validation of ambiguous calls. This approach greatly reduces the amount of manual labor involved in surveying landscapes when compared to traditional point counts or transect sampling. Nocturnal and cryptic species are widely understudied, not just in South India but all over the world.^45^ The amount of training data used to refine the detectors is the main reason why our current accuracy levels were at 56.3%; we need more annotated acoustic data from the tropics to ensure better detector accuracy. However, with the growing interest in PAM, and with hobbyist birders collecting acoustic recordings from species, this gap could be addressed. PAM can also be an efficient way to expand our acoustic datasets in data-deficient or data-sparse regions, to facilitate better acoustic models for species identification. Passive acoustic monitoring can therefore be a powerful tool to monitor biodiversity, especially over large spatial scales, in areas with low data coverage, and in areas with high biodiversity. The findings from such studies are important for conservation efforts by providing credible information to aid major policy changes, and can potentially lead the way for cryptic or rare species to be regionally discovered.^46^

## Author contributions

AG - Data analysis, Writing ; SCS - Data collection, Idea development, Reviewing, Writing; AJ - Reviewing; AS - Idea development, Reviewing, Funding; HK - Data collection, Idea development, Reviewing, Writing

## Acknowledgments

AG is a 12th-grade high school student at Chirec International School, planning on pursuing a career in the Environment. He would like to thank his Biology teacher, Dr. Sunitha Nagaraj, for her continued support and guidance in learning about the subject of ecology. We would like to Nthank our funders TIFRH (Department of Atomic Energy, Government of India, Project Identification No. RTI4007), and DBT (Ramalingaswami Re-entry Fellowship BT/RLF/Re-entry/07/2022 to AS). Thank you to Dr. Vijay Ramesh of Cornell University for providing us audio recorders, the Forest Department of Telangana for permits (RC NO. 11853/2017/WL-2), Dr. Gopalakrishna R., Viral Joshi, Sriram Reddy for expert assistance with identifying ambiguous calls, and Anirudh Singh for help with collating the migratory status of birds.

