## Supplementary Table 1 for "A Case Study: Using Passive Acoustic Monitoring Devices In A Growing Urban Landscape To Monitor Avian Diversity"

**Supplementary material**

*Supplementary Table 1*- Table of species misidentified[32]. Confidence of BirdNET detections and explanatory notes are shown above. Rows in brown were detected with a confidence above 0.95.

| **Type** | **Common name** | **Scientific name** | **Confidence of detector** | **Notes** |
|---|---|---|---|---|
| Waterfowl | Ruddy Shelduck | *Tadorna ferruginea* | 0.8762 | Dog |
| | Gadwall | *Mareca strepera* | 0.8048 | Frog |
| | Green-winged Teal | *Anas crecca* | 0.9679 | Not present |
| Grouse, Quail, and Allies | Gray Junglefowl | *Gallus sonneratii* | 0.8926 | Dog |
| | Black Francolin | *Francolinus francolinus* | 0.9206 | Never been reported in Telangana before |
| Pigeons and Doves | Rock Pigeon | *Columba livia* | 0.8209 | Noise |
| | Eurasian Collared-Dove | *Streptopelia decaocto* | 0.8656 | Plane, car horns |
| | Asian Emerald Dove | *Chalcophaps indica* | 0.7572 | Truck |
| Cuckoos | Banded Bay Cuckoo | *Cacomantis sonneratii* | 0.9709 | Red-wattled Lapwing |
| | Gray-bellied Cuckoo | *Cacomantis passerinus* | 0.7693 | A lapwing |
| | Fork-tailed Drongo-Cuckoo | *Surniculus dicruroides* | 0.7883 | Asian Koel |
| | Indian Cuckoo | *Cuculus micropterus* | 0.804 | Not present |
| Nightjars | Jerdon's Nightjar | *Caprimulgus atripennis* | 0.9279 | Noise |
| Rails, Gallinules, and Allies | Water Rail | *Rallus aquaticus* | 0.8873 | Never been reported in Telangana before |
| | Eurasian Coot | *Fulica atra* | 0.9866 | Truck horns |
| | Watercock | *Gallicrex cinerea* | 0.872 | Noise |
| Cranes | Common Crane | *Grus grus* | 0.9692 | Crickets |
| Shorebirds | Pied Avocet | *Recurvirostra avosetta* | 0.9136 | Indian Thick-knee |
| | Black-bellied Plover | *Pluvialis squatarola* | 0.8801 | A lapwing |
| | Common Ringed Plover | *Charadrius hiaticula* | 0.9421 | Indian Thick-knee |
| | Little Ringed Plover | *Thinornis dubius* | 0.8186 | Red-wattled Lapwing |
| | Gray-headed Lapwing | *Vanellus cinereus* | 0.8631 | Red-wattled Lapwing |
| | Whimbrel | *Numenius phaeopus* | 0.8788 | Vehicle horn |
| | Eurasian Curlew | *Numenius arquata* | 0.8341 | Dog |

| **Type** | **Common name** | **Scientific name** | **Confidence of detector** | **Notes** |
|---|---|---|---|---|
| | Black-tailed Godwit | *Limosa limosa* | 0.9936 | Red-wattled Lapwing |
| | Common Snipe | *Gallinago gallinago* | 0.9075 | Not present |
| | Common Sandpiper | *Actitis hypoleucos* | 0.9811 | Lesser Whistling-Duck in flight |
| | Common Redshank | *Tringa totanus* | 0.7767 | Red-wattled Lapwing |
| | Common Greenshank | *Tringa nebularia* | 0.8967 | Not present |
| | Dunlin | *Calidris alpina* | 0.9274 | Vehicle noise |
| | Small Buttonquail | *Turnix sylvaticus* | 0.9734 | Not present |
| Gulls, Terns, and Skimmers | Little Tern | *Sternula albifrons* | 0.89 | Red-wattled Lapwing |
| | Whiskered Tern | *Chlidonias hybrida* | 0.8446 | Long-tailed Shrike |
| | Common Tern | *Sterna hirundo* | 0.852 | Red-wattled Lapwing |
| | Sandwich Tern | *Thalasseus sandvicensis* | 0.9503 | Never been reported in Telangana before |
| Grebes | Little Grebe | *Tachybaptus ruficollis* | 0.7998 | Not present |
| Vultures, Hawks, and Allies | Osprey | *Pandion haliaetus* | 0.9068 | Common Myna |
| | Booted Eagle | *Hieraaetus pennatus* | 0.9694 | Indian Thick-knee |
| | Black Kite | *Milvus migrans* | 0.7939 | Indian Thick-knee |
| | Common Buzzard | *Buteo buteo* | 0.7894 | A lapwing |
| Owls | Indian Scops-Owl | *Otus bakkamoena* | 0.9655 | Dog |
| | Brown Boobok | *Ninox scutulata* | 0.7743 | Dog |
| Hoopoes | Eurasian Hoopoe | *Upupa epops* | 0.9635 | Vehicle horn |
| Falcons and Caracaras | Eurasian Kestrel | *Falco tinnunculus* | 0.7721 | Shikra mating call |
| | Peregrine Falcon | *Falco peregrinus* | 0.9608 | Red-wattled Lapwing |
| Larks | Crested Lark | *Galerida cristata* | 0.8502 | Never been reported in Telangana before |
| Cisticolas and Allies | Zitting Cisticola | *Cisticola juncidis* | 0.8269 | Not present |
| Martins and Swallows | Bank Swallow | *Riparia riparia* | 0.9611 | insect |
| | Western House-Martin | *Delichon urbicum* | 0.8829 | insect |
| Leaf Warblers | Hume's Warbler | *Phylloscopus humei* | 0.8035 | Not present |
| | Tickell's Leaf Warbler | *Phylloscopus affinis* | 0.8178 | Noise |
| | Common Chiffchaff | *Phylloscopus collybita* | 0.7632 | Indian Robin |

| **Type** | **Common name** | **Scientific name** | **Confidence of detector** | **Notes** |
|---|---|---|---|---|
| | Green Warbler | *Phylloscopus nitidus* | 0.763 | Indian Robin |
| Tree-Babblers, Scimitar-Babblers, and Allies | Indian Scimitar-Babbler | *Pomatorhinus horsfieldii* | 0.9644 | Abiotic sound |
| Flowerpeckers | Thick-billed Flowerpecker | *Pachyglossa agilis* | 0.7535 | Not present |
| Weavers and Allies | Baya Weaver | *Ploceus philippinus* | 0.8583 | Not present |
| Wagtails and Pipits | White Wagtail | *Motacilla alba* | 0.9451 | insect |
| | Paddyfield Pipit | *Anthus rufulus* | 0.9693 | insect |
| Finches, Euphonias, and Allies | Common Rosefinch | *Carpodacus erythrinus* | 0.8076 | Indian Robin |
